# A novel method of testing the antimicrobial potentials of *Exiguobacterium aurantiacum, Lysinibacillus boronitolerans* and *Bacillus megaterium*

**DOI:** 10.1101/2023.12.06.570382

**Authors:** Radhika Jain, Mugdha Belwalkar, Varsha Shukla, Anushree Lokur, Mayuri Rege

## Abstract

Microorganisms can be an effective source for antimicrobial compounds including peptides that provide an evolutionary advantage to them in specific niches. Prior studies have noted the antimicrobial properties of *Exiguobacterium aurantiacum, Lysinibacillus boronitolerans* and *Bacillus megaterium* but they used time- and resource-intensive procedures to prepare the extracts (2, 10). We wanted to explore a more cost-effective, time-effective and simpler method of testing the antimicrobial potential of bacterial extracts. Our extraction protocol is a one-step incubation of the organism in the solvent overnight, followed by a standard well diffusion based assay to test its antimicrobial effect. We tested this method with three cultures isolated from the premises of Mumbai railway stations: *E. aurantiacum, L. boronitolerans* and *Bacillus megaterium*. These extracts were tested against human pathogens *Staphylococcus aureus* and *Enterococcus faecalis. E. aurantiacum, L. boronitolerans* and *B. megaterium* inhibited *S. aureus* by 9.17 ± 0.42, 9.25 ± 0.48 and 10.50 ± 1.5 mm, respectively, and *E. faecalis* by 15.79 ± 2.25, 14.21 ± 0.88 and 17.5 ± 2.89 mm, respectively. This approach is therefore suggested to be effective for screening the antimicrobial activity of these bacteria.

## 1.0 Introduction and Background Information

Since ancient times, humans have faced disease caused by microorganisms (10). From pathogens such as *Helicobacter pylori* causing stomach infections over 100,000 years ago (1) to one of the most common diseases today, the leading cause of death in infants worldwide, Pneumonia, caused by the bacterium *Streptococcus pneumoniae* (16). Due to the potency of these infections, research investigating potential cures has been active since the 19th century (12). The first organism with antimicrobial activity was discovered in 1929 by Alexander Fleming - *Penicillium notatum*, which was found to produce Penicillin, a molecule that possessed the ability to inhibit bacterial growth (8). The search for antibiotics has intensified in the last few decades due to the growing number of antibiotic resistant microorganisms. Various natural sources like plants, actinomycetes, bacteria and fungi produce compounds that inhibit growth of human pathogens. But one of the major hurdles in the discovery of new antibiotics is a prolonged screening process (14).

Current methods to screen the antimicrobial potential of bacteria can be expensive and time-consuming. Common practices include the use of centrifugation, compound fractionation, materials such as Nutrient Broth, Lysogeny Broth and more (2, 10). Whilst these methodologies often provide accurate and precise results, they are also hard to replicate and scale as the apparatus required is costly. This is where the need for a simpler method of testing the antimicrobial activity of bacteria, that can be carried out using materials readily available in even school labs, arises.

In this study, the focus was placed on three bacteria that may have antimicrobial properties, namely *Exiguobacterium aurantiacum, Lysinibacillus boronitolerans* and *Bacillus megaterium* (6, 7, 9). Hence, the scope of research was narrowed to finding a novel method (that is potentially more cost-effective, time-effective and simpler) of testing the antimicrobial potentials of *E. aurantiacum, L. boronitolerans* and *B. megaterium* against common pathogens *Staphylococcus aureus* and *Enterococcus faecalis*.

### 2.0 Materials

The materials used were: Diethyl ether (Catalogue Number 00617, Research-Lab Fine Chem Industries, INDIA), Spinwin Microcentrifuge Tubes (Code 500020, Tarsons), Micropipette of capacity 100-1000 μL (Catalogue Number 380/08, Superfit), Micropipette of capacity 1-100 μL (Catalogue Number LHC37111016, Superfit), Toothpicks (Local Brand), 90 mm Nutrient Agar plates (item MP001, HiMedia Laboratories Pvt. Ltd., MH, INDIA), 100 mm Mueller-Hinton Agar plates (item MP173C, HiMedia Laboratories Pvt. Ltd., MH, INDIA), Sterilised swabs (Neptune Labs), 5 mm diameter cork borer (Neptune Labs), Forceps (Neptune Labs), 100-1000 μL micropipette tips (Neptune Labs), and 1-100 μL micropipette tips (Neptune Labs).

## 3.0 Methods

### 3.1 Collection and Isolation of Samples

The samples of *E. aurantiacum, L. boronitolerans* and *B. megaterium* were collected from local train stations in Mumbai, India, namely Chhatrapati Shivaji Maharaj Terminus (CSMT), Dadar, Dombivli and Thane stations (Fig. 1). Samples were picked up using sterile swabs, in accordance with standard sampling protocols (5). They were then streaked on Nutrient Agar plates and incubated overnight at 35°C. The grown cultures were identified using MALDI-TOF identification (see Additional Files 1, 2 and 3).

**Figure 1:**
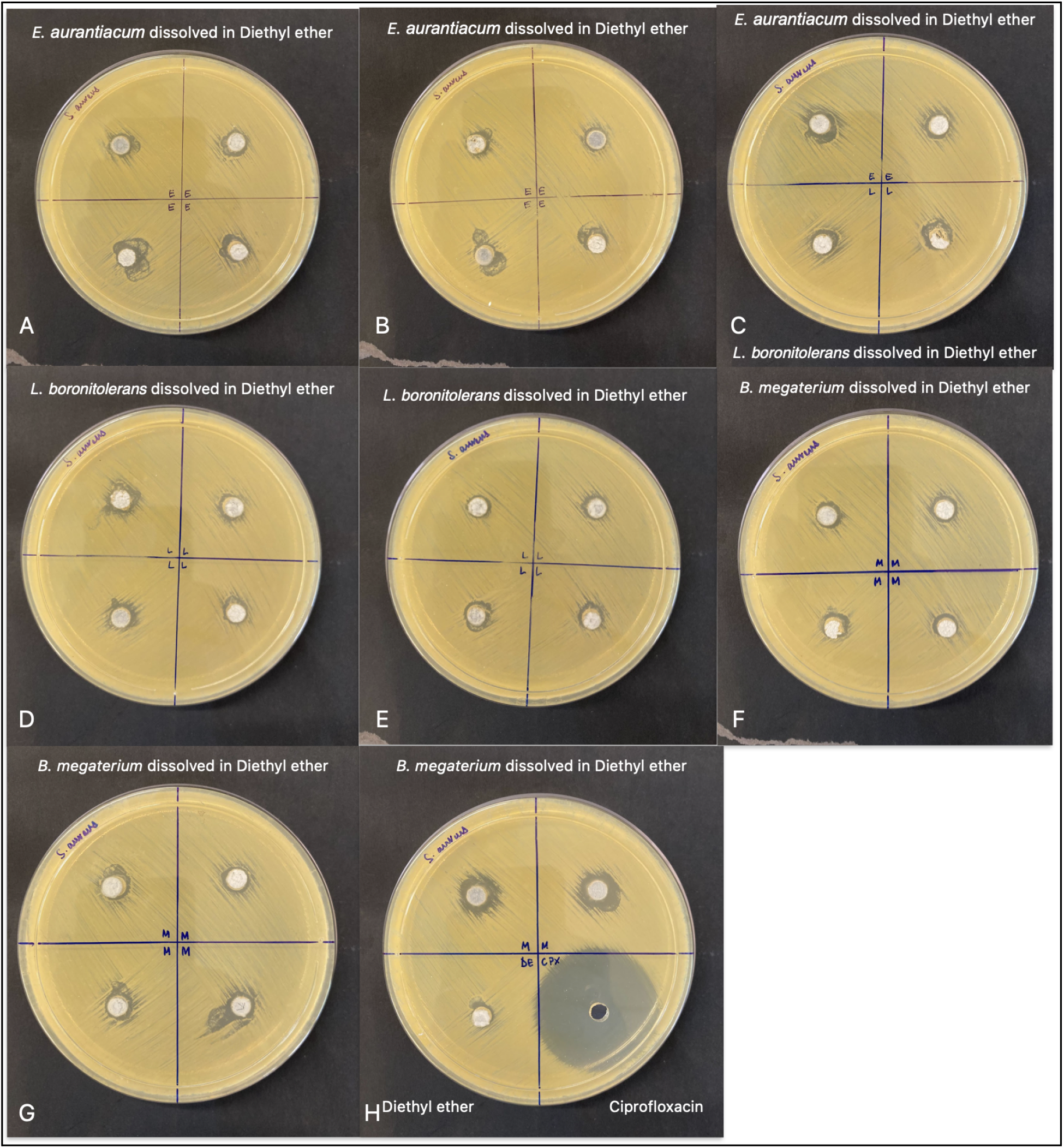
Testing the novel extraction method for antimicrobial activity against *S. aureus*. Standard well diffusion assays were performed to measure the zone of inhibition with bacterial extracts of *E. aurantiacum* (A-B and C top row), *L. boronitolerans* (C bottom row and D-E) and *B. megaterium* (F-H top row) against *S. aureus*. Negative control with only the solvent Diethyl ether and positive control with 1 mg/ml of Ciprofloxacin were also performed (H bottom row).

### 3.2 Source of Test Organisms

The strain of *S. aureus* used was identified using MALDI-TOF identification (see Additional File 4) and the strain of *E. faecalis* used was a typed strain.

### 3.3 Preparation of Solvent

Once the organisms had grown, one colony of each was picked up (using a toothpick) and dissolved in 2 ml of Diethyl ether in Eppendorf tubes. These tubes were placed in an incubator set at 30°C overnight. Diethyl ether was chosen as when dissolved in it, the bacterial isolates had exhibited antimicrobial activity in preliminary experiments. Preliminary experiments were also conducted with Ethyl acetate and Dichloromethane as solvents, but no antimicrobial properties were shown with these. An incubation temperature of 30°C was selected due to Diethyl ether’s low boiling point of 34.5°C (13).

### 3.4 Screening of Isolates for Antimicrobial Activity

The standard method of well diffusion was used for this study (3). Before choosing well diffusion as the final methodology, disk diffusion was also tested. However, it was found that well diffusion was far more effective due to the raised capacity for the solution.

One colony of the test culture was swabbed over the entire surface of the 100 mm Mueller-Hinton agar plate using a sterile swab stick. The plate was then rotated 90° and this was repeated for the second time. This was done twice more. The swabbing procedure was performed once more using another colony of the test culture. Four wells were made on each plate using a cork borer of diameter 5 mm and were filled with 120 μL of the solutions taken from the Eppendorf tubes using a micropipette of capacity 1-100 μL. Plates were incubated at 35°C for 24 hours and the zones of inhibition were examined in mm (10).

Negative controls of 120 μL of just the solvent (Diethyl ether) and positive controls of 120 μL of 1 mg/ml Ciprofloxacin were used to create points of comparison.

### 4.0 Results

Using the novel method, we tested one-step incubated extracts of three isolates (*E. aurantiacum, L. boronitolerans* and *B. megaterium*) for antimicrobial activity against two different strains known to be human pathogens, namely *S. aureus* and *E. faecalis*. The objective of this experiment was to investigate whether this efficient method of preparing bacterial extracts that hold antimicrobial properties can display zones of inhibition. After 24 hours in the incubator, the zone of inhibition (ZOI) around the wells (Figure 1 and 2) were recorded in mm for each trial and later averaged, as can be seen in Tables 1 and 2.

**Table 1:** Zones of inhibitions measured for all *S. aureus* conditions after 24 hours in the incubator.

| Treatment Type | Petri Dish Number | Zone of Inhibition (mm) | Average Zone of Inhibition (mm) | Mean Zone of Inhibition (mm) | Standard Deviation (± mm) |
| --- | --- | --- | --- | --- | --- |
| Positive Control (Ciprofloxacin) | 8 | 40.0 | 40.0 | 40.00 | - |
| Negative Control (Diethyl ether) | 8 | 0.0 | 0.0 | 0.00 | - |
| *Exiguobacterium aurantiacum* dissolved in Diethyl ether | 1 | 9.5, 9.0, 10.0, 8.5 | 9.25 | 9.17 | 0.42 |
|  | 2 | 9.0, 9.5, 9.5, 9.0 | 9.25 |  |  |
|  | 3 | 9.0, 9.0 | 9 |  |  |
| *Lysinibacillus boronitolerans* dissolved in Diethyl ether | 3 | 9.0, 10.0 | 9.5 | 9.25 | 0.48 |
|  | 4 | 10.0, 9.5, 9.0, 9.0 | 9.375 |  |  |
|  | 5 | 8.5, 9.0, 9.0, 9.0 | 8.875 |  |  |
| *Bacillus megaterium* dissolved in Diethyl ether | 6 | 9.0, 10.0, 9.0, 9.0 | 9.25 | 10.50 | 1.5 |
|  | 7 | 9.0, 10.0, 10.0, 9.0 | 9.5 |  |  |
|  | 8 | 13.0, 12.5 | 12.75 |  |  |

**Table 2:**
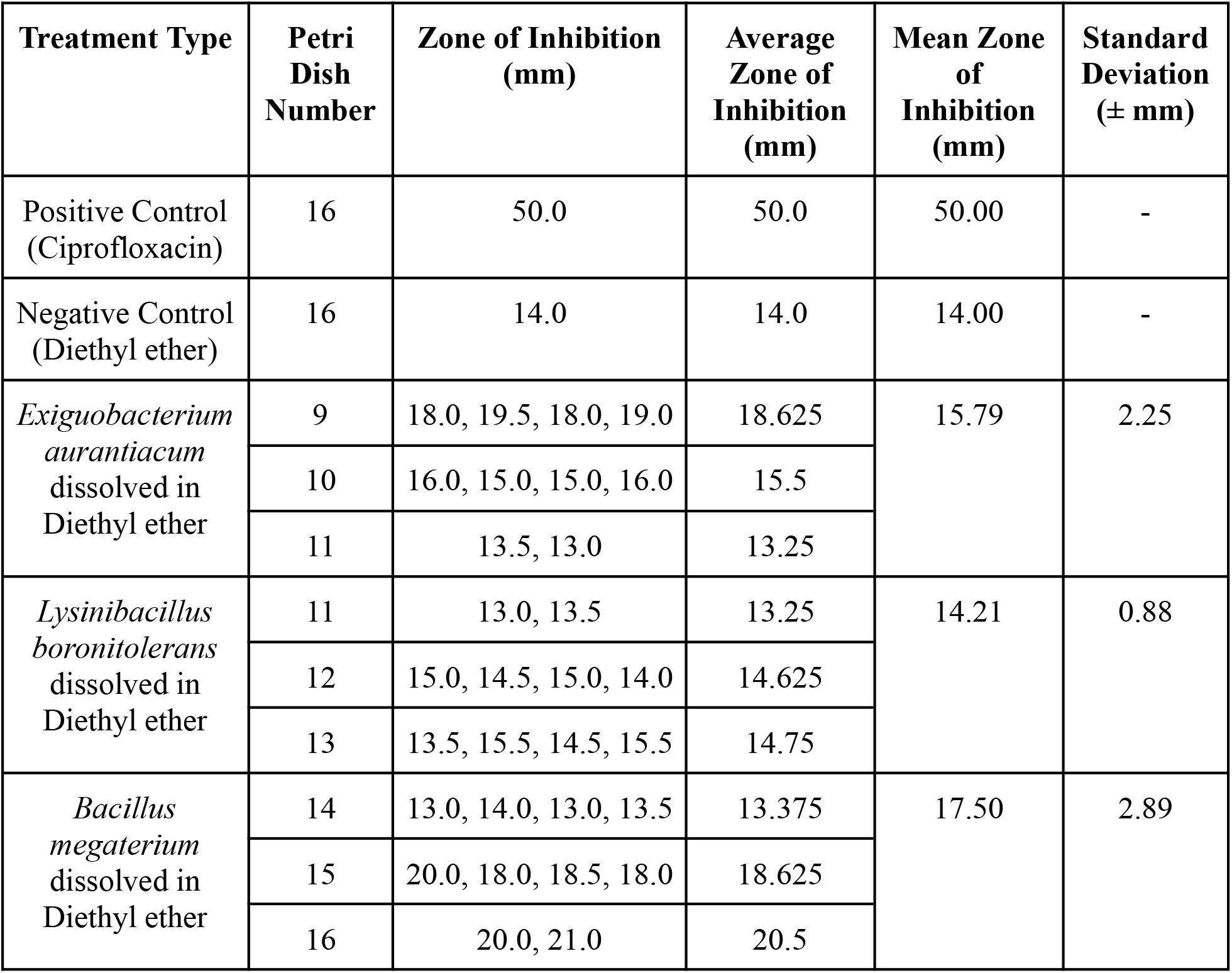
Zones of inhibitions measured for all *E. faecalis* conditions after 24 hours in the incubator.

**Figure 2:**
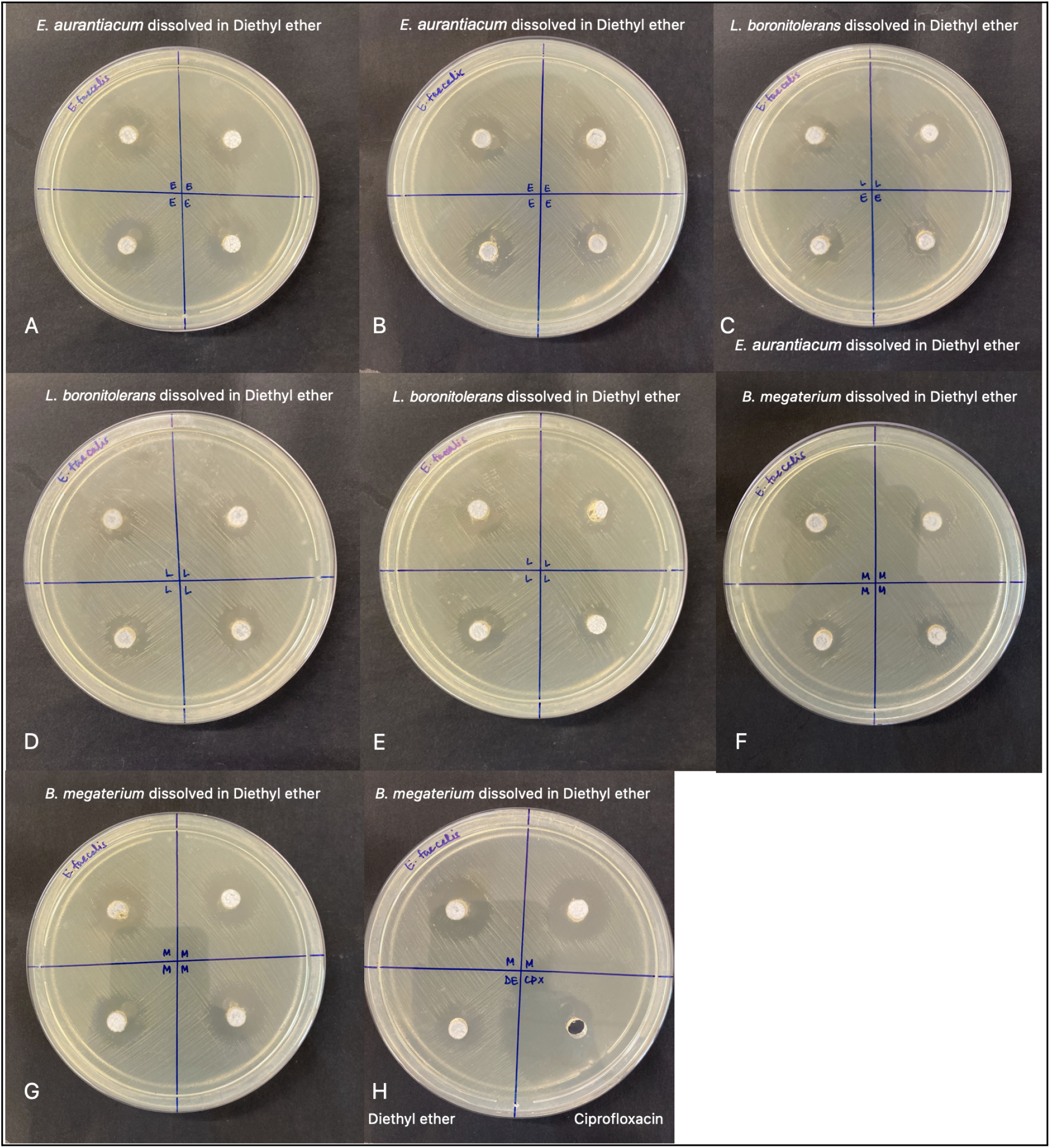
Testing the novel extraction method for antimicrobial activity against *E. faecalis*. Standard well diffusion assays were performed to measure the zone of inhibition with bacterial extracts of *E. aurantiacum* (A-B and C top row), *L. boronitolerans* (C bottom row and D-E) and *B. megaterium* (F-H top row) against *S. aureus*. Negative control with only the solvent Diethyl ether and positive control with 1 mg/ml of Ciprofloxacin were also performed (H bottom row).

Using only the solvent diethyl ether as a negative control, a 14 mm zone was observed against *E. faecalis* and no zone was observed against *S. aureus*. In contrast, large ZOI were observed against *S. aureus* and *E. faecalis* (of 40 and 50 mm respectively) with the positive control ciprofloxacin, as expected.

When tested against *S. aureus*, the one-step incubated extracts of *E. aurantiacum, L. boronitolerans* and *B. megaterium* all showed a ~10 fold greater inhibition compared to the negative control. Specifically, extracts of *E. aurantiacum* showed an average ZOI of 9.17 ± 0.42 mm, extracts of *L. boronitolerans* ZOI of 9.25 ± 0.48 mm and extracts of *B. megaterium* ZOI of 10.50 ± 1.50 mm. In *S. aureus*, the negative control (Diethyl ether) produced no measurable zone.

When tested against *E. faecalis*, extracts of *E. aurantiacum* showed an average ZOI of 15.79 ± 2.25 mm, extracts of *L. boronitolerans* ZOI of 14.21 ± 0.88 mm and extracts of *B. megaterium* ZOI of 17.50 ± 2.89 mm. As the negative control (Diethyl ether alone) gave a ZOI of 14 mm, the extract of *B. megaterium* showed the largest ZOI against this pathogen.

Overall, *B. megaterium* was seen to produce the largest zones of inhibition against both *S. aureus* and *E. faecalis* (10.50 ± 1.50 mm and 17.50 ± 2.89 mm respectively). *E. aurantiacum* produced the smallest zone out of the three bacteria against *S. aureus*, with an average zone of 9.17 ± 0.42 mm.

## 5.0 Discussion

The objective of this study was to find a cost-effective, time-efficient and simpler method of creating bacterial extracts to test their antimicrobial properties. The methodology developed is simple and functional, as zones of inhibition larger than the negative control were produced by most bacterial extracts against the two test pathogens. The new method was tested using microorganisms such as *Exiguobacterium aurantiacum, Lysinibacillus boronitolerans, Bacillus megaterium*, that are known to have antimicrobial properties against gram positive and gram negative bacteria (2, 7, 9). However prior studies used technically challenging methods to extract antimicrobial substances from these bacteria. Our method may provide a simpler, faster and quick screening method to test antimicrobial properties of bacteria.

However, some limitations should be noted. Only one trial of the positive and negative control were performed for each test pathogen, precluding analysis using a statistical test such as a *t*-test. Additionally, the zones produced by *L. boronitolerans* dissolved in Diethyl ether against *E. faecalis* were slightly larger than that of the negative control (14.21 mm and 14.00 mm, respectively). Another limitation of this methodology is that Diethyl ether, the solvent in use, is highly volatile, flammable and therefore, slightly inefficient to use (4). Larger amounts of the extracts than needed had to be prepared as half the quantities evaporated in the overnight incubation period. Additionally, Diethyl ether forms a white precipitate on plastic surfaces such as the plastic petri dishes used (can be seen in the wells of the plate where the Diethyl ether was pipetted, see Figures 1 and 2), but does not form precipitates on glass petri dishes (Additional File 5). Lastly, there was little variance between the zones produced by *E. aurantiacum* and *L. boronitolerans* against *S. aureus*. This could be explained by similarities in mechanism of action, which is yet to be determined.

This work is relevant to the field of antibiotic screening, as it provides an efficient method for preliminary testing for antimicrobial properties in bacteria. There are also several extensions of this research. To find the compound that is exhibiting these antimicrobial properties, the prepared solvents could be separated and tested for activity using bioautography (15). Compounds that inhibit the growth of test pathogens could be identified using a mass spectrometric analysis. Additionally, more trials and variations of this research could be performed in order to further investigate the relationship between these solvents and other antibiotics. For example, as seen in petri dishes 8 and 16, *B. megaterium* produced larger zones in the presence of Ciprofloxacin in comparison to when it was tested alone (petri dishes 6, 7, 14 and 15). This suggests the possibility of a synergistic relationship between Ciprofloxacin and the active compound in the *B. megaterium* solution and can be further explored through more experiments.

## 6.0 Conclusion

On average, *B. megaterium* extracts produced the largest zone of inhibition against both *S. aureus* and *E. faecalis*. Further experiments and statistical analysis are required in order to identify the active compounds exhibiting antimicrobial properties, but this methodology seems promising as an efficient preliminary method of testing for antimicrobial properties of bacteria.

## Supporting information

Supplemental files Ruia college

## 7.0 Acknowledgements

We would like to thank members of the Department of Microbiology at Ramnarain Ruia Autonomous College Mumbai, Mr Omkar Nadkarni and Ms Supriya Sawant at BIS, Mumbai and my parents for their guidance and support throughout this research project.

## 8.0 Author Contributions

Roles based on PLOS ONE standards (11).

Radhika Jain: Investigation and Original Draft Preparation

Radhika Jain, Mayuri Rege, Mugdha Belwalkar: Conceptualization, Data Curation, Formal Analysis, Methodology

Mugdha Belwalkar, Mayuri Rege: Supervision, Review & Editing

Mayuri Rege, Mugdha Belwalkar, Varsha Shukla, Anushree Lokur: Funding Acquisition and Resources

