## Supplemental files Ruia college for "A novel method of testing the antimicrobial potentials of *Exiguobacterium aurantiacum, Lysinibacillus boronitolerans* and *Bacillus megaterium*"

HiMedia Laboratories Pvt Ltd

Plot No:C40, Road 21Y, Wagle Industrial Estate, Thane West, Maharashtra 400604,India




### Autof ms1000 Identification Report

#### Basic Information

Sample Name: MTI00040_Sample 4

Sample Spot: F6

Sample Description: T3_TNA_NA_4

Operator: Tanuja Patil

Generated Time: 02/17/2023 17:15

Identification Result: Exiguobacterium aurantiacum 9.566

#### Mass Spectrum

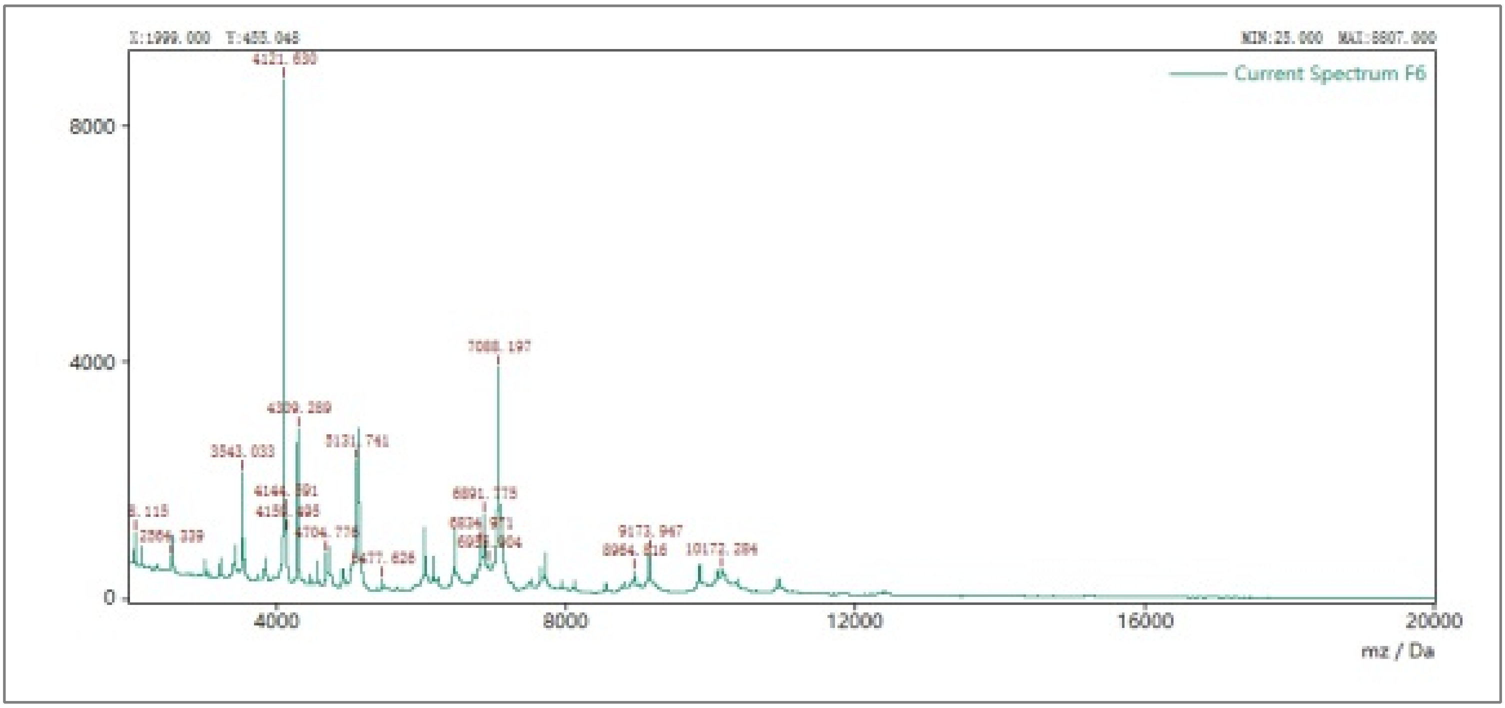


#### Detailed Results

| NO. | Result | Score |
|---|---|---|
| 1 | Exiguobacterium aurantiacum | 9.566 |

##### Interpretation of score system

| Score Range | Result Interpretation |
|---|---|
| 9.500 to 10.000 | Credible upto species & subspecies level |
| 9.000 to 9.500 | Credible upto species level |
| 6.000 to 9.000 | Credible upto genus level |
| 0.0 to 6.000 | Unreliable* |

*: May need further investigation

Done By
(Microbiologist/ Sr. Microbiologist)

Checked By
(Asst/Dy/Manager - Microbiology)

Approved By
(Vice President/ Sr.Vice President- Microbiology)

HIMEDIA
FOR LIFE IS PRECIOUS

HiMedia Laboratories Pvt Ltd

Plot No:C40, Road 21Y, Wagle Industrial Estate, Thane West, Maharashtra 400604,India



### Autof ms1000 Identification Report

#### Basic Information

Sample Name: MSN00063_TPN82127113_sample027

Sample Spot: F2

Sample Description: T4_TNA_MA_5

Operator: Tanuja Patil

Generated Time: 05/06/2023 18:45

Identification Result: Lysinibacillus boronitolerans 9.281

#### Mass Spectrum

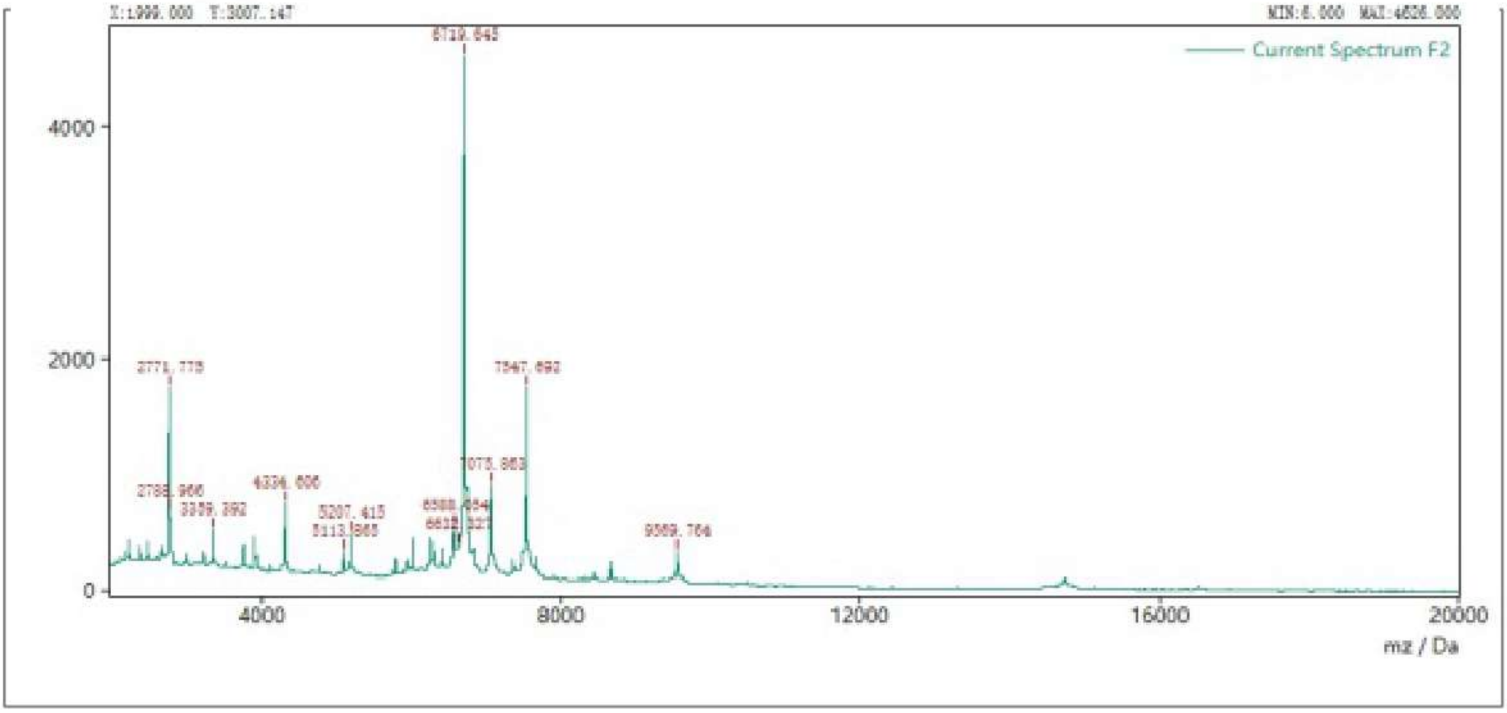


#### Detailed Results

| NO. | Result | Score |
|---|---|---|
| 1 | Lysinibacillus boronitolerans | 9.281 |
| 2 | Lysinibacillus boronitolerans | 9.159 |
| 3 | Paenibacillus chibensis | 4.882 |
| 4 | Bordetella petrii | 4.753 |
| 5 | Penicillium oxalicum | 4.407 |
| 6 | Penicillium oxalicum | 4.218 |
| 7 | Penicillium oxalicum | 4.149 |
| 8 | Chryseobacterium gleum | 4.135 |
| 9 | Candida inconspicua | 4.127 |
| 10 | Aspergillus niger | 4.113 |

### HiMedia Laboratories Pvt Ltd

Additional File 3

Plot No:C40, Road 21Y, Wagle Industrial Estate, Thane West, Maharashtra 400604,India


#### Autof ms1000 Identification Report

##### Basic Information

Sample Name: MISN00043_TPN82127113_Sample006

Sample Spot: A7

Sample Description: T2_CSMT_1

Operator: Tanuja Patil

Generated Time: 03/10/2023 11:09

Identification Result: Bacillus megaterium 9.313

##### Mass Spectrum

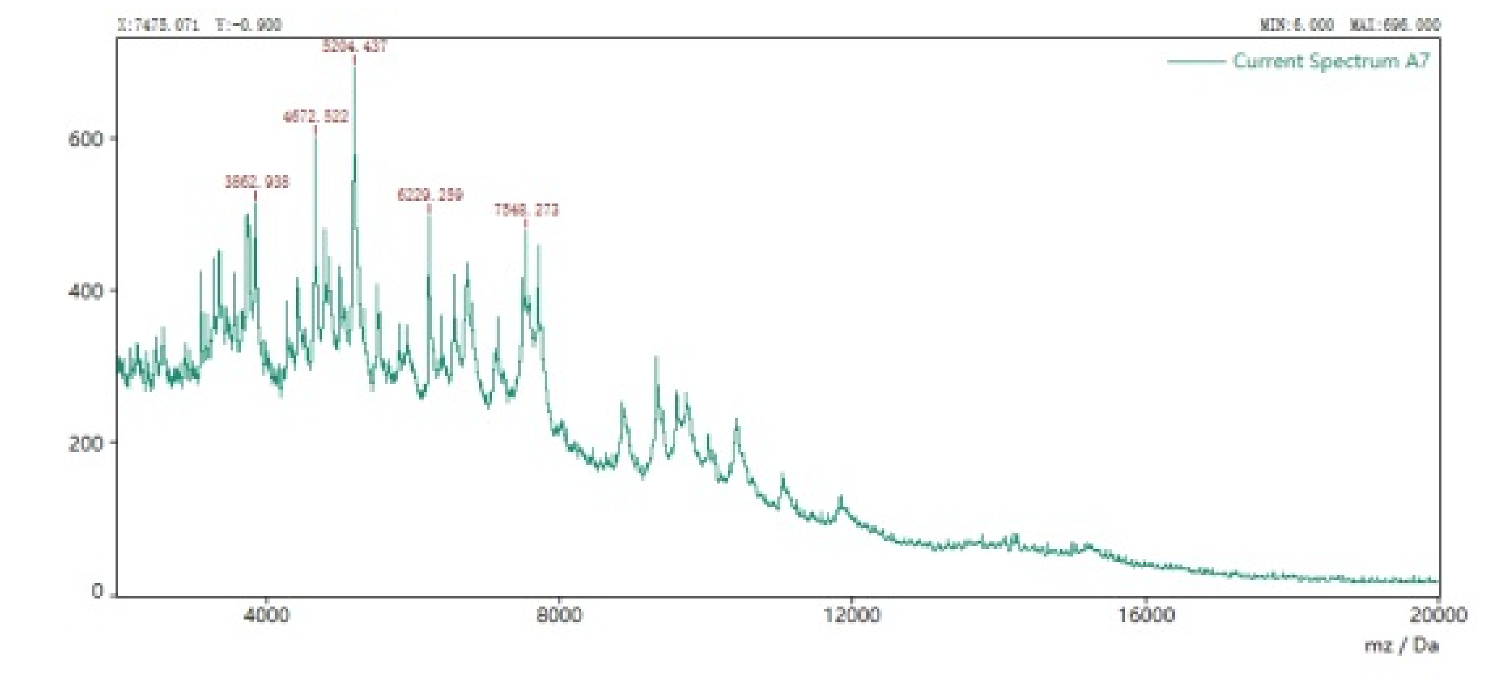


##### Detailed Results

| NO. | Result | Score |
|---|---|---|
| 1 | Bacillus megaterium | 9.313 |

###### Interpretation of score system

| Score Range | Result Interpretation |
|---|---|
| 9.500 to 10.000 | Credible upto species & subspecies level |
| 9.000 to 9.500 | Credible upto species level |
| 6.000 to 9.000 | Credible upto genus level |
| 0.0 to 6.000 | Unreliable* |

*: May need further investigation

Done By
(Microbiologist/ Sr. Microbiologist)

Checked By
(Asst/Dy/Manager - Microbiology)

Approved By
(Vice President/ Sr.Vice President- Microbiology)

Additional File 4

HiMedia Laboratories Pvt Ltd
Plot No:C40, Road 21Y, Wagle Industrial Estate, Thane West, Maharashtra 400604,India
ph:-022-61471919 Website: www.himedialabs.com

### Autof ms1000 Identification Report

#### Basic Information

Sample Name: MISN00087_TPN82127114_sample004

Sample Spot: D8

Sample Description: S.a

Operator: Tanuja Patil

Generated Time: 07/24/2023 15:34

Identification Result: Staphylococcus aureus 9.297

#### Mass Spectrum

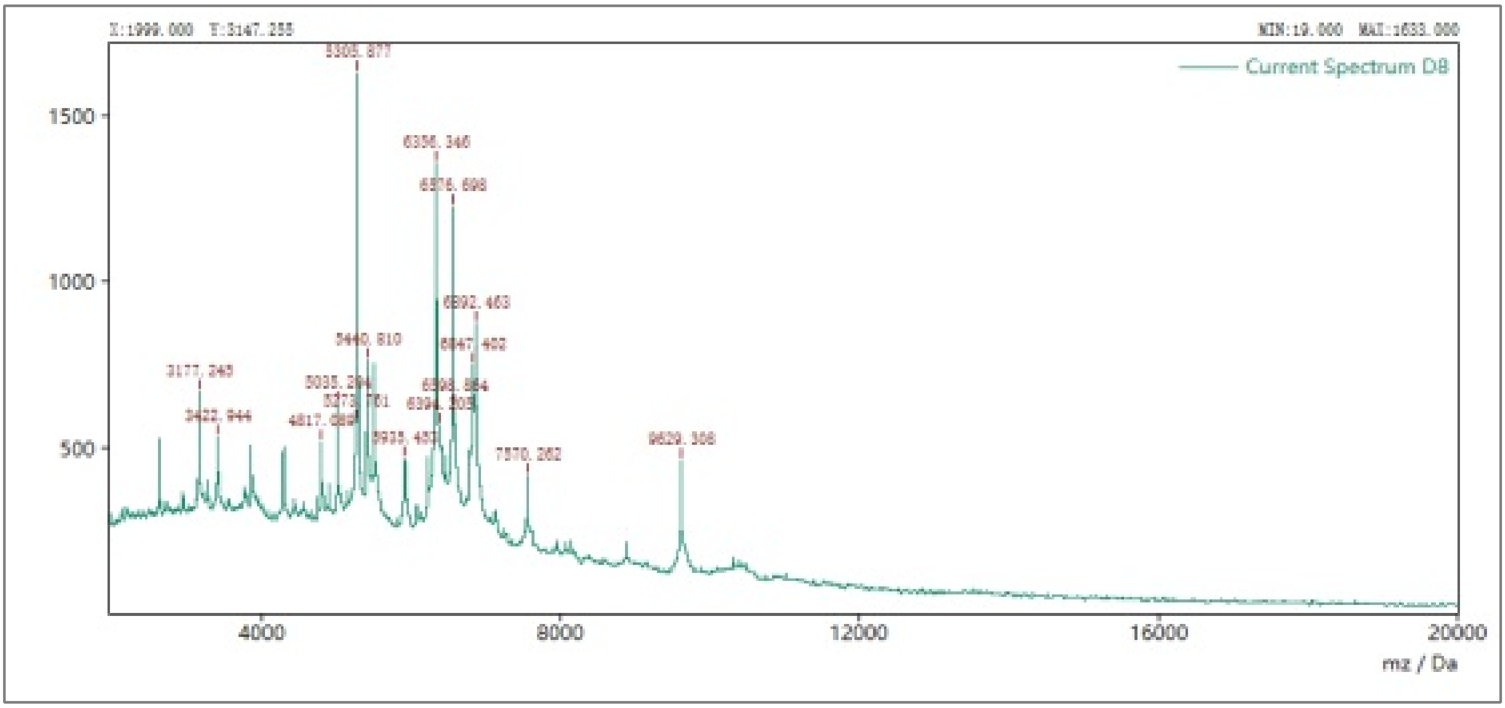


#### Detailed Results

| NO. | Result | Score |
|---|---|---|
| 1 | Staphylococcus aureus | 9.297 |
| 2 | Staphylococcus aureus | 9.135 |
| 3 | Staphylococcus aureus | 9.079 |
| 4 | Staphylococcus aureus | 9.057 |
| 5 | Staphylococcus aureus | 8.923 |
| 6 | Staphylococcus aureus | 8.837 |
| 7 | Staphylococcus aureus | 8.808 |
| 8 | Staphylococcus aureus | 8.793 |
| 9 | Staphylococcus aureus | 8.710 |
| 10 | Staphylococcus aureus | 8.601 |

### Additional File 5

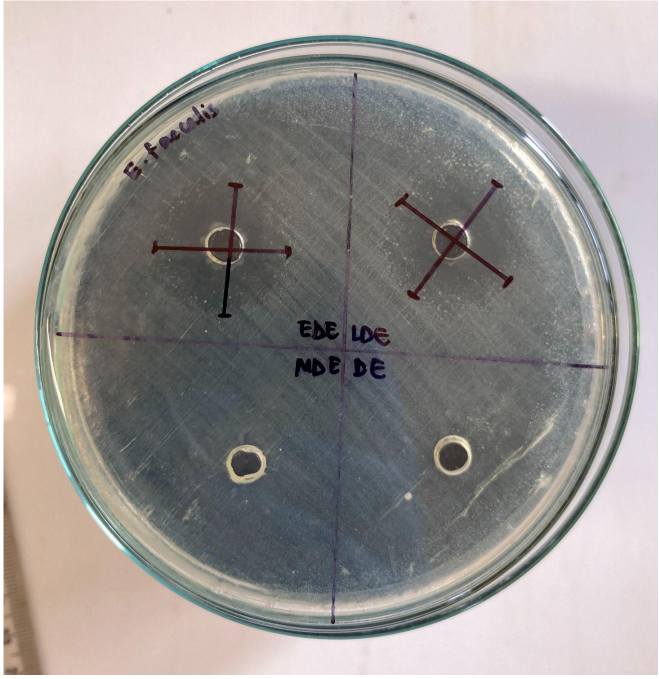


*Figure 3: Experiment done using a glass petri dish, where there is no white precipitate formed by the Diethyl ether*
